# Joint modelling of diffusion MRI and microscopy

**DOI:** 10.1101/563809

**Authors:** Amy FD Howard, Jeroen Mollink, Michiel Kleinnijenhuis, Menuka Pallebage-Gamarallage, Matteo Bastiani, Michiel Cottaar, Karla L Miller, Saad Jbabdi

## Abstract

The combination of diffusion MRI with microscopy provides unique opportunities to study microstructural features of tissue, particularly when acquired in the same sample. Microscopy is frequently used to validate diffusion MRI microstructure models, addressing the indirect nature of dMRI signals. Typically, these modalities are analysed separately, and microscopy is taken as a gold standard against which dMRI-derived parameters are validated. Here we propose an alternative approach in which we combine diffusion MRI and microscopy data obtained from the same tissue sample to drive a single, joint model. This simultaneous analysis allows us to take advantage of the breadth of information provided by complementary data acquired from different modalities. By applying this framework to a spherical-deconvolution analysis, we are able to overcome a known degeneracy between fibre dispersion and radial diffusion. Spherical-deconvolution based approaches typically estimate a global fibre response function to determine the fibre orientation distribution in each voxel. However, the assumption of a ‘brain-wide’ fibre response function may be challenged if the diffusion characteristics of white matter vary across the brain. Using a generative joint dMRI-histology model, we demonstrate that the fibre response function is dependent on local anatomy, and that current spherical-deconvolution based models may be overestimating dispersion and underestimating the number of distinct fibre populations per voxel.

## 1. Introduction

Diffusion MRI (dMRI) is routinely used to study white matter microstructure and connectivity *in vivo* and non-invasively [1, 2, 3]. dMRI microstructure models relate variations in the MR signal to microstructural features of interest. Such inference requires biophysical modelling of both the tissue architecture and diffusion process. Although many dMRI models have been proposed, few have been rigorously validated [4, 5], and the link between the observed diffusion signal and the underlying white matter microstructure remains controversial [6, 7].

Microscopy is often considered a gold standard technique for the validation of dMRI models. Crucially, microscopy tends to resolve a specific structure of interest (e.g. histological staining of astrocytes or polarised light imaging of myelinated axons) and thus typically provides specificity that is not guaranteed by MRI. In a typical validation study the dMRI and microscopy data are analysed separately, then dMRI-derived tissue parameters (e.g. fibre orientation, myelin density or axon diameter) are compared to microscopy equivalents which are taken to be the ground truth [8, 9, 10, 11]. This is possible due to the complementary nature of the data: both modalities provide information about the same tissue parameters of interest, but each observe them through a different lens. However, by analysing the data separately (rather than simultaneously), such paradigms may not be exploiting the multimodal data to its full potential.

Here we suggest an alternative, data-fusion framework in which we combine dMRI and microscopy data from the same tissue sample into a single joint model. A joint model may be advantageous in three respects. Firstly, by considering both datasets simultaneously, we have access to additional, complementary information about the tissue microstructure and may be able to accurately determine tissue parameters that are currently unobtainable from the diffusion signal alone. A secondary benefit of the data-fusion framework is that the joint model considers both dMRI and microscopy to be informative of the ‘true’ underlying microstructure, but also that both have sources of uncertainty (Figure 1). Crucially, these are unique, modality-dependent sources of noise. Therefore, by using a data-fusion framework we can in theory obtain a higher-precision estimate of the underlying microstructure of interest. Finally, microscopy is typically 2D and may only be sensitive to a subset of the tissue compartments (e.g. myelinated axons or astrocytes). For example, histological staining of the tissue (a gold standard microscopy technique) typically produces 2D images of thin tissue sections, where only the stained microstructure is easily visualised. Thus, the information provided by microscopy only partially informs on the tissue microstructure. The joint model can overcome this limitation by considering the microscopy as a soft constraint on the model, as opposed to a hard constraint or ground truth in post-hoc validation. This framework is inspired by a similar data-fusion approach [12] which demonstrated improved brain connectivity analysis when complementary 3T and 7T dMRI data was analysed jointly rather than separately.

**Figure 1:**
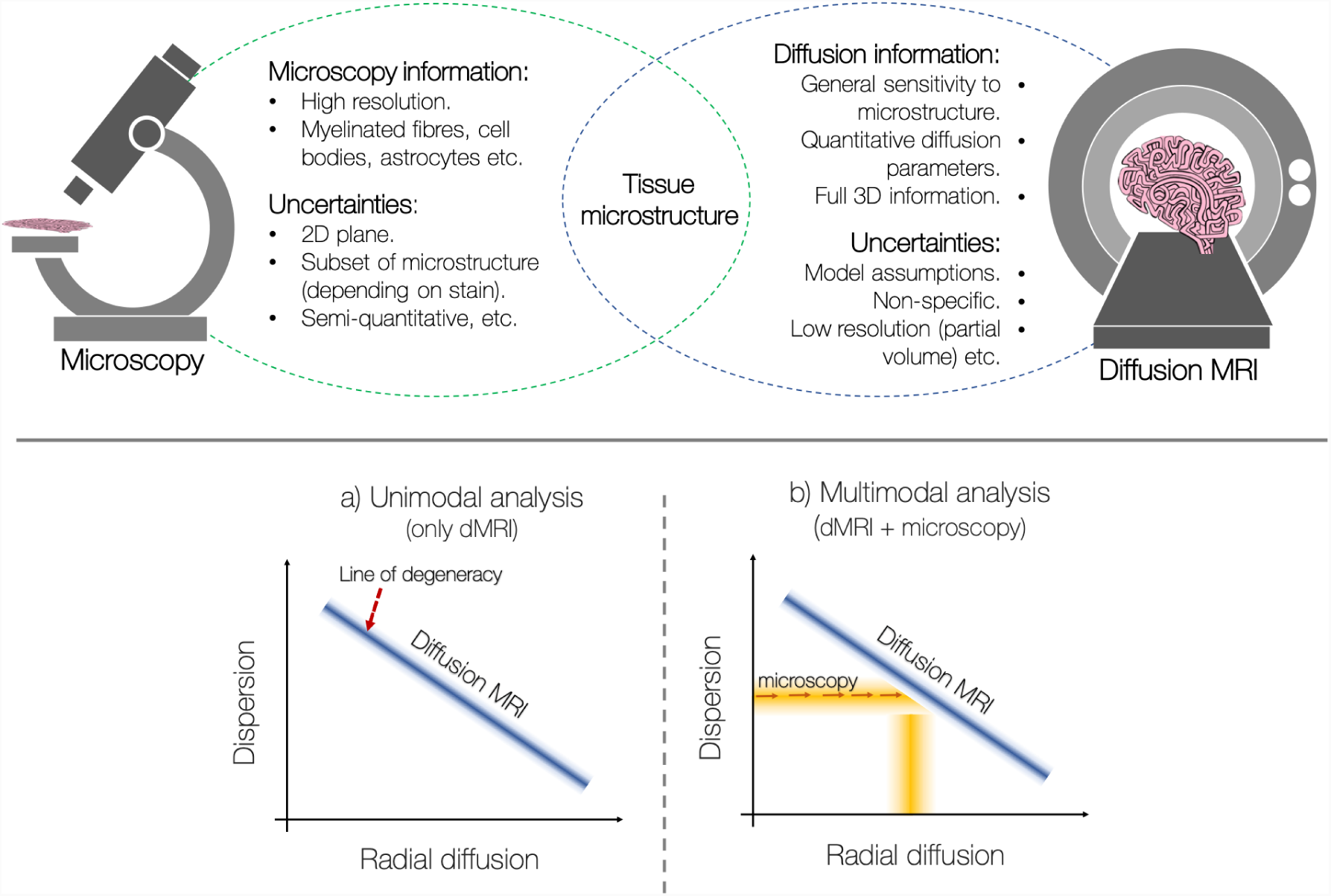
Top: Microscopy data can provide highly detailed information about specific microstructural features of the tissue at sub-micrometre resolutions. In comparison, the diffusion MRI signal is an indirect measure of the same microstructure of interest. Due to the complementary information provided by the two modalities, we propose a data-fusion framework which simultaneously analyses dMRI and microscopy data from the same tissue sample to drive a single, joint model. Bottom: (a) A highly dispersed fibre population with low radial diffusivity may produce the same diffusion MRI signal as a single, coherently ordered fibre population with high radial diffusivity. The blue line labelled ‘Diffusion MRI’ represents a simplistic but graphical representation of this degeneracy. (b) Microscopy can estimate and, in the joint analysis, constrain the fibre dispersion in each MRI voxel to overcome the degeneracy; the dMRI data can now provide accurate estimates of radial diffusion. Critically, as the microscopy acts as a soft constraint (rather than fixing the dispersion), both the dMRI and the microscopy data simultaneously inform on the fibre dispersion. Information about the radial diffusion comes from dMRI data alone. The shading represents noise in the data and uncertainty in the parameter estimates.

This report considers one example of how dMRI-microscopy data-fusion allows us to extract tissue parameters which are difficult to obtain from the dMRI data alone. Here we aim to separate out fibre orientation dispersion from radial diffusion i.e. the apparent diffusion coefficient perpendicular to the direction of the fibre. As is illustrated in Figure 1 (bottom), a highly dispersed fibre population with low radial diffusivity may produce the same dMRI signal as a single, coherently oriented fibre population with high radial diffusion. This degeneracy is commonly overcome by assuming one can identify a region with a single, coherently oriented fibre population which is then used to define a global radial diffusivity. Alternatively, promising developments by Lasič *et al.* [13], Szczepankiewicz *et al.* [14], and more recently by Cottaar *et al.* [15] demonstrate that, by combining linear and spherical diffusion tensor encoding, it is possible to disentangle microscopic diffusion anisotropy from orientation dispersion. Nonetheless, current analyses of conventional dMRI data (based on single diffusion encoding and which do not assume global diffusivities) are unable to distinguish these two distinct fibre configurations.

Microscopy can be used to determine the fibre orientations at a much finer spatial resolution (typically ∼ micrometer or sub-micrometer per pixel) and so can estimate the fibre dispersion in each dMRI voxel. However, microscopy alone is typically uninformative of the diffusion properties of the tissue. In a joint dMRI-microscopy model (as illustrated in Figure 1, bottom), once the amount of within-voxel fibre dispersion is constrained, dMRI can accurately estimate the radial diffusion. In this manner a joint model should be able to overcome the degeneracy and separate out fibre dispersion from radial diffusion. This could provide insights into how these two parameters vary across the brain in both health and disease.

There are many ways to formulate a joint model. In this report we focus on one approach based on constrained spherical deconvolution (CSD, [16]). As a popular data-driven analysis, CSD estimates the underlying fibre orientations from a dMRI dataset. In CSD, the diffusion signal is considered to be the convolution of the fibre orientation distribution (FOD) with a fibre response function (FRF). Here the FRF describes the diffusion signal from a single, coherently oriented fibre population. In the model, the FRF is first estimated empirically (typically from voxels with large fractional anisotropy [16, 17, 18, 19]) and subsequently used as a global deconvolution kernel to determine the underlying FOD. CSD is thus based around two main assumptions: that it is possible to identify a region which contains a single, non-dispersed fibre population, and that the FRF estimated from this region holds globally. In other words, it is possible to estimate a valid ‘brain-wide’ FRF. These assumptions may be challenged if we consider there to be non-zero orientation dispersion in typical MRI voxels, and the diffusion characteristics of white matter to be dependent on microstructural properties such as axonal diameter, packing and myelination that can vary across the brain [20, 21]. Thus, the estimation of a brain-wide fibre response function could be unreliable and has been shown to produce spurious results [22, 17] when poorly estimated. Alongside the recent and complimentary publication by Schilling *et al.* [23] this report actively demonstrates the extent of FRF variation across the brain.

In this work we combine dMRI and histology data from the same human tissue sample to investigate the diffusion properties of white matter under conditions where multiple modalities are informative of the ‘true’ fibre configurations. To highlight the benefits of the data-fusion framework, this report considers a joint model which is based on non-parametric CSD. We show that by including histology data into the joint model we are able to over-come the degeneracy of fibre dispersion and radial diffusion. As such, we can simultaneously estimate the diffusion profile of a single fibre bundle (the FRF) and the underlying fibre orientation distribution (the FOD) on a voxel-by-voxel basis. In our results we investigate how the FRF changes across several white matter regions, and consider the implications this may have on both the reliability of CSD-based analyses and our understanding of the white matter microstructure.

## 2. Methods

This section will first describe the principles of CSD, and how these were developed into our joint modelling approach (c.f. *Joint modelling*). As a generative model which spans multiple modalities, the joint model error balances three terms (a dMRI-data fidelity term, a microscopy-data fidelity term and a complexity penalty) with two regularisation factors. Thus, *Simulations* describes how simulated data (with a known ground truth) was used to determine appropriate values for the two regularisation factors. Finally, *Postmortem data acquisition* provides details of the co-registered high-resolution dMRI (0.4mm isotropic) and myelin-stained histological data used in this study.

### 2.1 Constrained spherical deconvolution

In constrained spherical deconvolution (CSD) the diffusion-weighted MR signal attenuation, *S*, measured along an orientation parametrised in spherical coordinates by angles (*θ*_0_, *φ*_0_) is considered to be the convolution of the FOD with the single-fibre response function (FRF):

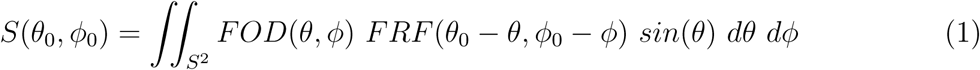

where both the FOD and FRF are defined on the unit sphere. The FRF describes the signal profile of a single coherently-oriented fibre bundle and both the FRF and the FOD can be expressed in terms of spherical harmonics [16].

After estimating an appropriate FRF, the FODs are reconstructed on a voxel-wise basis by deconvolving the FRF from the diffusion signal *S*. As the deconvolution is sensitive to noise, the spherical deconvolution is constrained to minimise physically impossible negative peaks in the reconstructed FOD [24]. To constrain the optimisation a modified Tikhonov regularisation method [25] is typically employed to iteratively penalise negative amplitudes on the FOD and update the FOD estimation.

This report will compare results from the standard CSD model (a deconvolution) with those from the novel joint modelling approach (a generative model). In the standard CSD model, processing was performed using the MRtrix3 package (www.mrtrix.org, [26]) and the FRF was determined empirically using Tournier’s approach [27]. Briefly, the Tournier algorithm iterates between response function estimation and CSD to determine the 300 most likely ‘single-fibre’ voxels from which the FRF is estimated.

### 2.2 Joint modelling

The principles of constrained spherical deconvolution were extended to enable joint estimation of the FRF and FOD for datasets with both dMRI and microscopy data. Although the joint modelling approach could be applied to any type of microscopy data from which we can extract fibre orientations, in this work we analyse histological images which have been stained for myelin and co-registered to 0.4 mm isotropic dMRI data (c.f. *2.4 Postmortem data acquisition*). Through structure tensor analysis [28] of histological images, the primary fibre orientation was estimated per pixel after which 1400 x 1400 orientations were combined into a frequency histogram to generate a 2D microscopy-derived FOD (*FOD*_2*D,micro*_) for a ‘superpixel’ comparable to the spatial resolution of dMRI. In the model, the microscopy-derived FOD acted as a soft constraint on both the fibre orientation and amount of dispersion in each MRI voxel. This allowed estimation of the diffusivities both parallel (*d*_*axial*_) and perpendicular (*d*_*radial*_) to the fibre.

The generative model is described in Figure 2. In the joint model, the FRF and FOD were combined to predict the dMRI signal, *S*, along each gradient direction, *k*(*θ*_0_, *φ*_0_), using equation (1). Additionally, the FOD was projected onto the 2D plane (*FOD*_2*D,joint*_) and re-normalised to fit the microscopy-derived *FOD*_2*D,micro*_. To retain simplicity and avoid overfitting, in the joint model the FRF was characterised by an axially-symmetric diffusion tensor, whilst the FOD was defined on a spherical harmonic basis set of order 6. The model thus contained 30 free parameters: 2 diffusivities (which determined the FRF) and 28 spherical harmonics coefficients (which determined the ODF).

**Figure 2:**
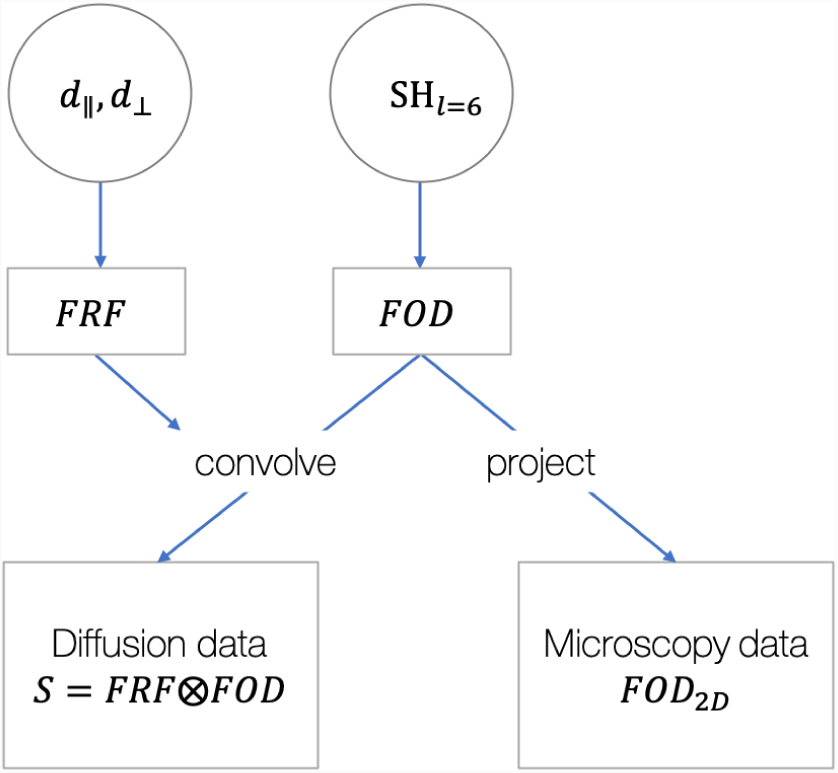
The generative joint model. Here the fibre response function (FRF) was considered an axially symmetric diffusion tensor characterised by diffusivities (*d*) parallel and perpendicular to the fibre. The fibre orientation distribution (FOD) was defined on a spherical harmonic basis set of order *l* = 6. In the joint model, the FRF and FOD were simultaneously fit to the diffusion data (through convolution) and the FOD projected onto a 2D plane for comparison with the microscopy-derived FOD.

The model parameters were fitted by minimising a cost function *E* made of three terms,

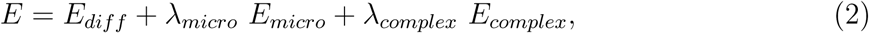

where *E*_*diff*_ describes the dMRI-data fidelity term, *E*_*micro*_ the microscopy-data fidelity term, and *E*_*complex*_ a complexity penalty. The predicted dMRI signal, *S*_*k*_, was compared to the dMRI data, *Y*_*k*_, using the mean squared error,

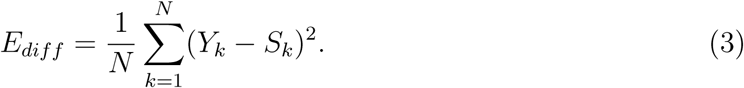

The projected FOD (*FOD*_2*D,joint*_) was compared to the microscopy-derived *FOD*_2*D,micro*_ using the symmetric Kullback-Leibler divergence *D*_*KL*_ [29],

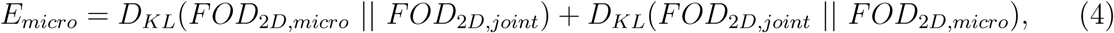

where the Kullback-Leibler divergence of two discrete probability distributions P and Q is defined as,

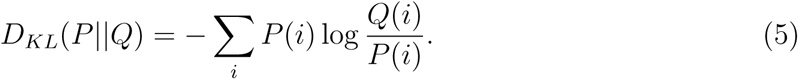

To minimise the presence of spurious peaks in the FOD, a third error term, the complexity penalty, penalised non-zero spherical harmonic coefficients which were not supported by the data. Here the complexity penalty minimised the L1-norm of the spherical harmonics coefficients, *x*, which describe the 3D FOD where,

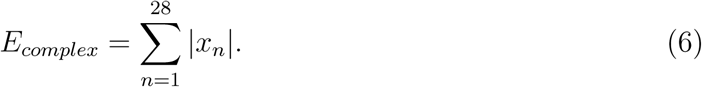

The three error terms were combined into a single cost function (Equation 2), with regularisation factors *λ*_*micro*_ and *λ*_*complex*_ respectively, and minimised using MATLAB’s non-linear solver *fmincon* [30]. Optimisation constraints ensured that the FOD, *d*_*axial*_ and *d*_*radial*_ were positive and that *d*_*axial*_ ≥ *d*_*radial*_. The joint model was initialised as follows: in each voxel, the spherical harmonics coefficients output from CSD [16] provided an initial estimation of the FOD and the FRF was initialised to *d*_*axial*_ = 0.25 *µm*^2^*/ms, d*_*radial*_ = 0.05 *µm*^2^*/ms*. MATLABs simulated annealing algorithm *simulannealbnd* [30] was used to generate three sets of starting parameters from the above initial conditions. The model was optimised for each set of starting parameters, and the solution with the lowest error chosen. With this procedure, the joint model was able to simultaneously optimise both the FRF and FOD on a voxel-by-voxel basis.

### 2.3 Simulations

This section will describe two sets of simulations which were used to determine appropriate regularisation factors *λ*_*micro*_ and *λ*_*complex*_. dMRI and histology data was simulated for a fairly simple single-fibre configuration. The first set of simulations was used to determine an appropriate value of *λ*_*micro*_ and did not include a complexity penalty (*λ*_*complex*_ = 0). We evaluated whether, through the inclusion of histology data (increasing *λ*_*micro*_), the joint model was able to separate fibre dispersion from radial diffusion by assessing how faithfully the model could quantify the fibre orientation dispersion. Here we included 3 parameters: the azimuth, *θ*, and inclination, *ϕ*, of the FOD peak, and the symmetric fibre dispersion around the peak, characterised by the orientation dispersion index (*ODI*, Equation 7). In the second set, *λ*_*micro*_ was fixed and a cross-validation approach was taken to optimise *λ*_*complex*_. Here spherical harmonics were used to describe the FOD and we assessed how the residual error of the model changed with respect to *λ*_*complex*_.

In both sets of simulations a dispersed FOD was approximated by a Watson distribution with concentration parameter *κ* [31, 32]. As proposed by Zhang et al. [32], an orientation dispersion index was defined as,

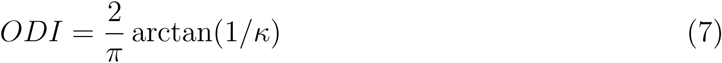

whereby the *ODI* varies from 0 (no dispersion) to 1 (isotropic dispersion). As above, the diffusion signal was calculated as the convolution of the FOD with an axially symmetric FRF (Equation 1) after which zero-mean Gaussian noise was added. In a similar manner to Sotiropoulos *et al.* [33], assuming *S*_0_ = 100, the SNR was defined as *SNR* = *S*_0_*/σ*_*noise*_. To simulate histology data, fibre orientations were sampled from the Watson distribution using rejection sampling, and then projected onto a 2D histological plane and combined into a frequency histogram. 1400 × 1400 samples were drawn from the Watson distribution to match the number of fibre orientations in each ‘superpixel’ of the real data. No additional noise was added to the histology data. Simulation parameters were fixed to values of *ODI* = 0.25, *d*_*axial*_ = 0.2 *µm*^2^*/ms, d*_*radial*_ = 0.1 *µm*^2^*/ms, SNR*_*dMRI*_ = 15.

We first determined an appropriate value for the weighting of the histology data *λ*_*micro*_, whilst setting *λ*_*complex*_ = 0. As we were primarily interested in how accurately the model could separate fibre dispersion from radial diffusion, a wrapper function was used to simplify the joint model such that the model contained only 3 free parameters: *ODI, d*_*axial*_ and *d*_*radial*_. To do this the fibre orientation was fixed to a specific direction, (*θ, φ*), after which the Watson-like FOD could be fully characterised by the *ODI* alone (rather than the 28 spherical harmonics of the normal joint model). dMRI and histology data was simulated for a single-fibre population oriented along (*θ, φ*) and a given *ODI*. The simplified joint model was fit to the simulated data using a Markov chain Monte Carlo (MCMC) method, Metropolis Hastings [34], to find optimal values of *ODI, d*_*axial*_ and *d*_*radial*_. *d*_*axial*_ and *d*_*radial*_ were initialised using the same parameters as for the real data (*d*_*axial*_ = 0.25 *µm*^2^*/ms, d*_*radial*_ = 0.05 *µm*^2^*/ms*) whilst the dispersion was initialised to *ODI* = 0.5. The optimisation was constrained such that the *ODI, d*_*axial*_ and *d*_*radial*_ were positive and *d*_*axial*_ ≥ *d*_*radial*_. It was preferable here to use MCMC for optimisation (instead of *fmincon* above) for two reasons. Firstly, the multiple MCMC samples identify combinations of parameters which produce an equally good fit. This allowed us to investigate both the precision and accuracy of the model. Secondly, as there were fewer model parameters to fit (2 diffusivities and the *ODI*, rather than the 30 free parameters above), the optimisation was better suited to MCMC methods. In these simulations of a single fibre population, the number of model parameters was greatly reduced as we fit a single-fibre FOD of symmetric dispersion, characterised by the *ODI*. In real data, we describe the FOD using spherical harmonics coefficients rather than the *ODI* of a Watson distribution, allowing for greater complexity in the FOD shape (e.g. multiple peaks with complex dispersion). The above procedure was first performed for a fibre population oriented along the histological plane (*φ* = 0) where the histological data was most informative of the ODF shape, and then repeated for various inclinations, *φ*.

For the second set of simulations, a cross-validation approach was used to determine an appropriate weighting of the model complexity penalty, *λ*_*complex*_. The complexity penalty aims to minimise spurious peaks in the FOD by penalising non-zero spherical harmonics which are unsupported by the data. If *λ*_*complex*_ were too low it would be ineffective, but if set too high, the spherical harmonic coefficients would be unable to accurately describe the FOD. The latter would be identified in a cross-validation study as it would result in a high error of the model with respect to a validation dataset. For the cross-validation study, data was simulated for a single-fibre population (as described above) and divided into training and validation datasets of equal proportions. That is, each set included half of the gradient directions, distributed fairly evenly across the sphere. For various *λ*_*complex*_, the model parameters *d*_*axial*_, *d*_*radial*_, and the spherical harmonic coefficients were fit to the training data, after which the out-of-sample residual error was calculated.

### 2.4 Postmortem data acquisition

This study analyses formerly obtained dMRI and histology data; for a detailed description of samples, the data acquisition and post processing see [10]. Briefly, postmortem brain tissues, which had been immersion fixed in formalin, were obtained from the Oxford Brain Bank, Nuffield Department of Clinical Neurosciences, University of Oxford, Oxford, UK. The postmortem tissues were from three male donors with no known neurological conditions who died of non-brain related disease. From each brain, a 5mm thick coronal section was extracted at the level of the anterior commissure to include various anatomical regions of interest: the corpus callosum, centrum semiovale, corticospinal tract, and cingulum bundle.

dMRI data was acquired on a 9.4T preclinical MRI scanner equipped with a 100 mm bore gradient insert (Varian Inc, CA, USA) and with a maximum gradient strength of 400 mT/m. At a spatial resolution of 0.4 mm isotropic, a diffusion-weighting of b = 5000 s/mm^2^ was achieved with a single-line spin-echo sequence (TE=29 ms, TR = 2.4s) over a total of 120 gradient directions. An additional eight images were collected with negligible diffusion weighting (b = 8 s/mm^2^).

After MRI scanning, the tissue was sectioned in half along the anterior-posterior direction. One half was processed for classic immunohistochemical staining of myelin and the other for polarised light imaging. As previous work [10] determined that the myelin staining provided better predictions of dispersion (those more consistent with dMRI data), this study focused solely on these histological images. For immunohistochemical staining, the tissue was paraffin embedded and sectioned at 6 *µ*m along the coronal plane. To visualise the myelin content, three sections of tissue were stained with antibodies against proteolipid protein (MCA839G; Bio-Rad; 1:1000) and imaged at a spatial resolution of 0.28 *µ*m/pixel. Structure tensor analysis [28, 35] of the histological images estimated the primary fibre orientation per pixel. 1400 × 1400 fibre orientations were then combined into a frequency histogram to generate a 2D histology-derived FOD at a spatial resolution comparable to the 0.4 mm dMRI data.

To allow for voxel-wise comparison of the multimodal data, the dMRI and histology were co-registered using a 2D registration based on a Modality Independent Neighbourhood Descriptor (MIND) [36] algorithm. Previous evaluation of the registered data [10] found especially good alignment in the corpus callosum facilitating robust voxel-wise comparisons. However, slight mis-alignment was apparent at tissue boundaries, particularly in specimen 3.

## 3. Results

### 3.1 Model validation

Using simulated data we first assessed whether the joint model was able to separate fibre dispersion from radial diffusion as intended. To determine an appropriate weighting of the histological data, *λ*_*micro*_, we examined the model specificity and accuracy when estimating *d*_*axial*_, *d*_*radial*_ and the *ODI*. Figure 3 considers a FOD oriented along the histological plane, where the histology is most informative. Figure 3a shows samples from fitting a simulated FOD using MCMC where the *ODI, d*_*axial*_ and *d*_*radial*_ were optimised simultaneously. With only dMRI data included in the model (Figure 3a, left), we found a clear degeneracy between fibre dispersion and radial diffusion as expected. A similar degeneracy exists between *d*_*axial*_ and the *ODI* which is not shown. In comparison, Figure 3a (right) depicts the same fit but with both histology and dMRI data included in the model. Here the histology data acted to constrain the within-voxel fibre dispersion. Even though the histology data was only informative of the fibre dispersion in 2D, the joint model was able to overcome the degeneracy and estimate both the fibre dispersion and radial diffusion simultaneously.

**Figure 3:**
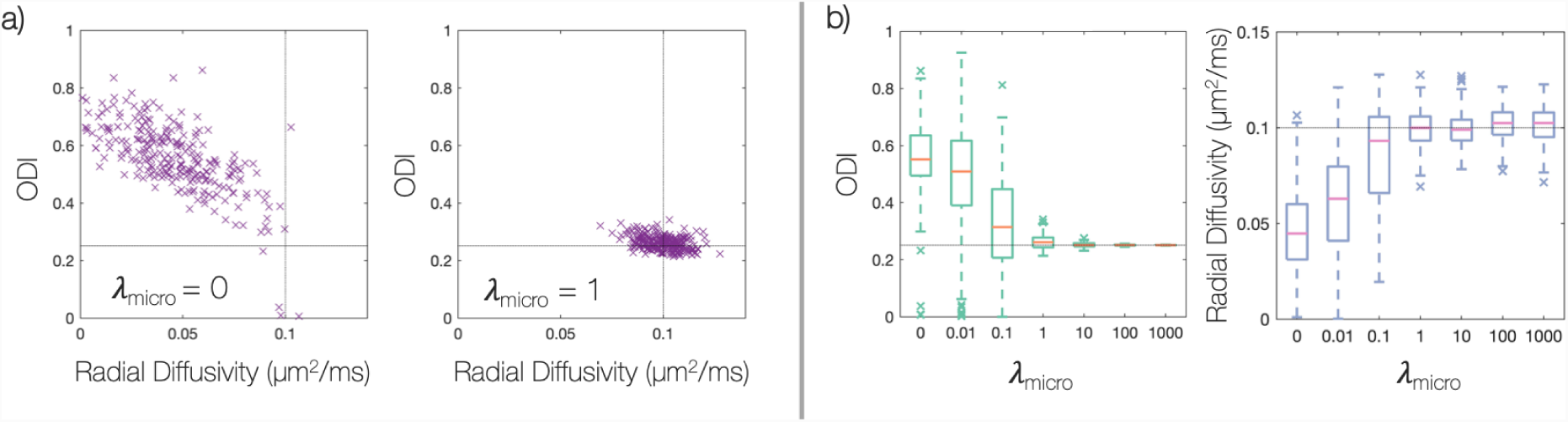
Overcoming the degeneracy between fibre dispersion and radial diffusion. Here, simulated data for a single-fibre FOD has been fit using MCMC. Note, the fibre orientation dispersion index (*ODI*) ranges from 0 (no dispersion) to 1 (isotropic dispersion) and the light grey lines represents the ground truth. Here we simulate data for a single fibre population oriented along the histological plane, where the histological data is most informative. a) Each point shows a sample from MCMC where the cost function of the joint model has been minimised with equally good fit. When the model only considers the dMRI data (left), we see a clear degeneracy as expected. However, the degeneracy may be overcome through the inclusion of histology data to the model (right). Part b) examines the effect of *λ*_*micro*_, the weighting of the histological data. When *λ*_*micro*_ = 1, the joint model was able to estimate both the *ODI* and radial diffusivity to a high degree of accuracy and precision.

In the model, the histologically-derived error was weighted by a regularisation factor *λ*_*micro*_; when *λ*_*micro*_ = 0, the histological data was excluded from the model, corresponding to the left part of 3a. Figure 3b shows how the accuracy of estimating *d*_*radial*_ and the *ODI* varies with respect to *λ*_*micro*_. When *λ*_*micro*_ = 1, we observe a break in the degeneracy between *d*_*radial*_ and the *ODI*, demonstrated by the increase in precision and accuracy.

In Figures 3a,b the FOD was oriented along the histological plane, such that the 2D histological data was highly informative of the FOD shape. In contrast, the histological data should be minimally informative for FODs oriented out of the histological plane. Figure 4 evaluates the performance of the model for fibre FODs with increasing inclination, *φ*, with respect to the histological plane. For *λ*_*micro*_ = 1, both *d*_*radial*_ and the *ODI* were estimated with good accuracy for fibre inclinations of up to 60 degrees. Accurate estimation of *d*_*radial*_ and the *ODI* for fibres of higher inclination required increasing values of *λ*_*micro*_. Finally, for an FOD oriented perpendicular to the histological plane, the degeneracy between the *ODI* and *d*_*radial*_ remained for all *λ*_*micro*_.

**Figure 4:**
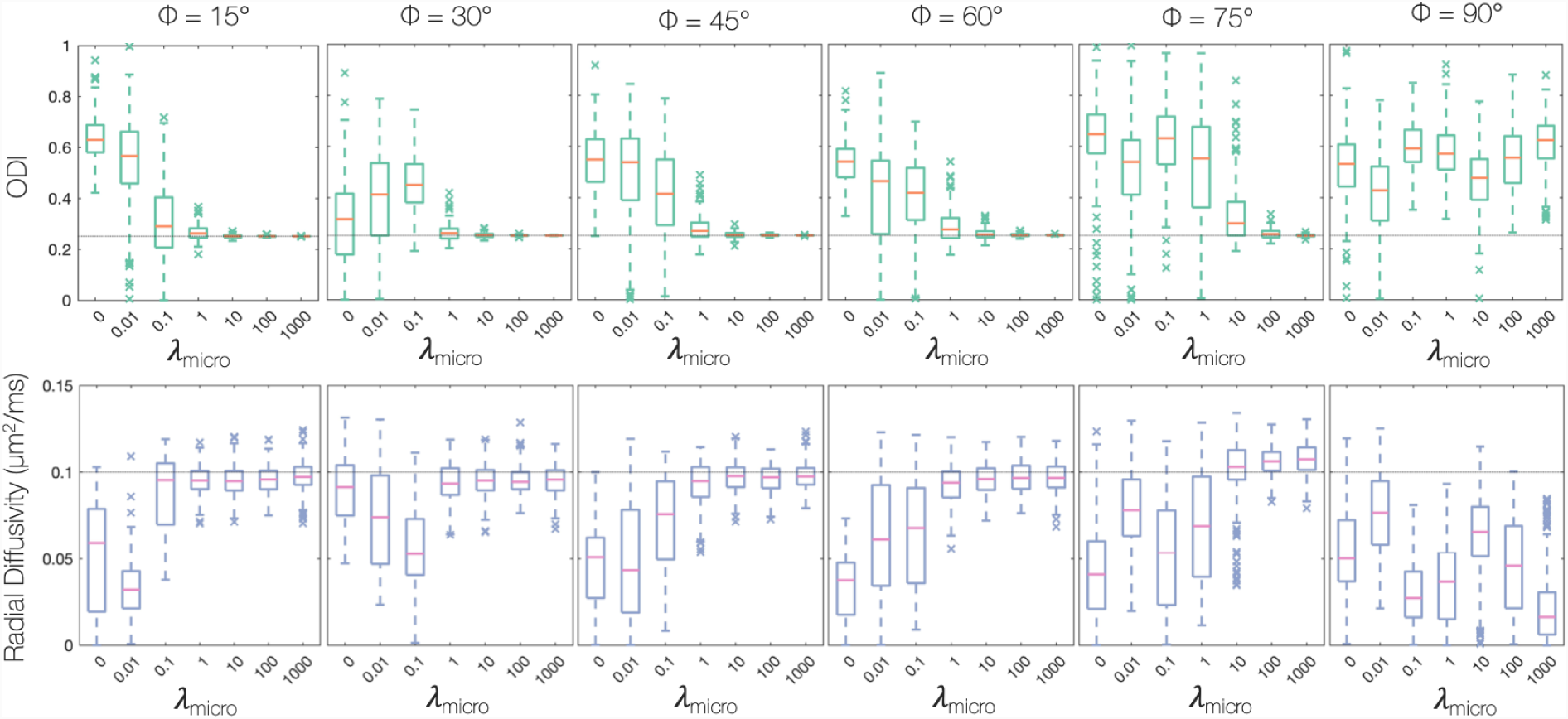
How the accuracy of the joint model, for a given *λ*_*micro*_, is affected by the FOD inclination with respect to the histological plane, *φ*. As in Figure 3, data was simulated for a single-fibre FOD and fit using MCMC. The light grey line represents ground truth values. When *λ*_*micro*_ = 1, the joint model was able to faithfully estimate both the *ODI* and radial diffusivity for fibres of inclinations up to 60^°^ out of the histological plane.

A high value of *λ*_*micro*_ may imbalance the microscopy data-fidelity term, *E*_*micro*_, with respect to the dMRI-data fidelity term, *E*_*diff*_, and result in a poor fit of the model to the dMRI data. Therefore, *λ*_*micro*_ = 1 was chosen for all further optimisations.

With *λ*_*micro*_ = 1, a procedure of cross-validation was used to determine the weighting of the complexity penalty, *λ*_*complex*_. Figure 5 plots the out-of-sample residual error with respect to *λ*_*complex*_. To remove spurious peaks from the FOD requires the highest complexity penalty supported by the data (i.e. one which does not detrimentally increase the residual error). Consequently, *λ*_*complex*_ = 0.01 was considered optimal.

**Figure 5:**
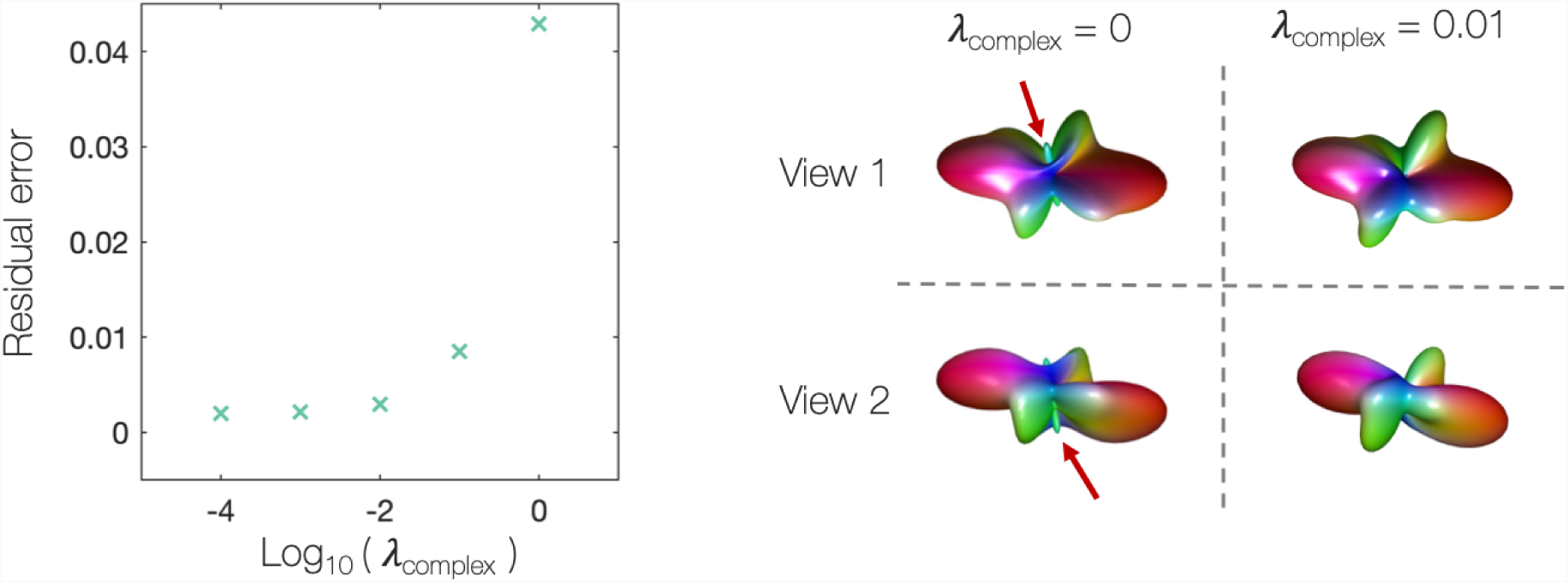
Determining *λ*_*complex*_, the weighting of the complexity penalty. A complexity penalty was added to prevent spurious peaks occurring in the FOD. Left: Having performed cross-validation on simulated data, the residual error of the out-of-sample data was estimated for *λ*_*complex*_ ranging from 1 ×10^-4^ to 1. *λ*_*complex*_ = 1 × 10^-2^ was chosen for all future optimisations due to the unsubstantial increase in the residual error. Right: An example voxel from the postmortem dataset. The voxel was optimised both without a complexity penalty (*λ*_*complex*_ = 0) and with (*λ*_*complex*_ = 0.01). We see how the complexity penalty successfully minimises the presence of a likely artefactual peak in the FOD, highlighted by the red arrows.

### 3.2 Residual error

Figure 6 compares the fit of the joint model and CSD to both the dMRI and histological data. For standard CSD, the FRF was estimated empirically from the most likely ‘single-fibre’ voxels using the Tournier algorithm [27]. Once estimated, the FRF was held constant across the sample and used as a deconvolution kernel to estimate, in each voxel, the ODF from the dMRI data. In comparison, the joint model acted as a forwards model, fitting to both the dMRI and histology data to estimate the FRF and ODF on a voxel-by-voxel basis. Figure 6 shows the mean absolute percentage error between the predicted diffusion signal (convolution of the ODF and FRF, Equation 1) and the dMRI data (top), and the Kullback-Leibler divergence (Equations 4, 5) between the histological data and the model FOD when projected into the histological plane (bottom).

**Figure 6:**
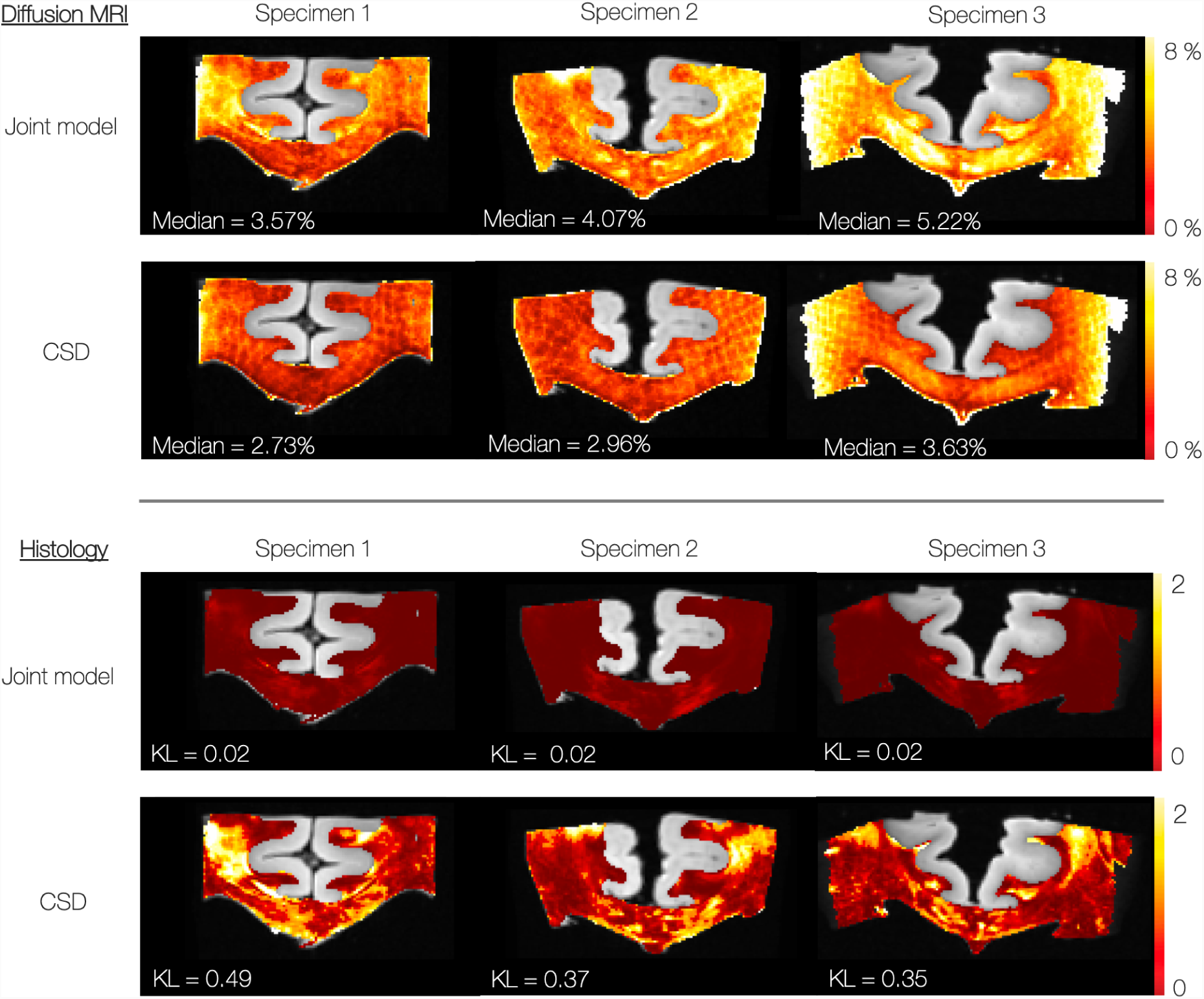
Top: The residual error (calculated as the mean absolute percentage error) of the model fit to the dMRI data only. Bottom: The error of the model with respect to the histological data, defined as the symmetric KullbackLeibler (KL) divergence. When KL = 0, the 2D-FODs are identical. Compared to CSD, there is a slight increase in the residual error of the joint model with respect to the dMRI data. This is to accommodate the highly improved fit to the histological data.

When compared to CSD, the joint model shows a slight increase in the dMRI-associated error which is concurrent with a largely improved fit to the histological data. This is as expected: CSD by definition minimises the error with respect to the dMRI data. In comparison, the joint model considers additional information about the ‘true’ fibre dispersion (from the histology data) to avoid overfitting.

### 3.3 Variations in axial and radial diffusivities

Standard CSD methods define a global FRF and thus assume that the diffusion characteristics of the tissue remain constant across the sample. In contrast, the joint model overcomes the degeneracy between fibre dispersion and radial diffusion to estimate the FRF on a voxel-by-voxel basis. In the joint model the FRF is considered an axially symmetric diffusion tensor described by *d*_*axial*_ and *d*_*radial*_. So, the joint model can estimate how the axial and radial diffusivities (and thus the FRF) vary voxel-wise across the sample.

Figure 7 shows considerable variation in the estimated axial and radial diffusivities across all three specimens. Here we should again note the less accurate registration of dMRI and histology data in specimen 3. Previous work [10] demonstrates the mis-alignment of the tissue around the grey matter tissue boundaries of specimen 3 as well as the more robust registration across the corpus callosum. Values of axial and radial diffusivity outside of the corpus callosum in specimen 3 should therefore be viewed with a sceptical eye. Nonetheless, in Figure 7 (top) we generally see a similar pattern of variation in both axial and radial diffusivities, with particularly low values of diffusivity often found in the corpus callosum.

**Figure 7:**
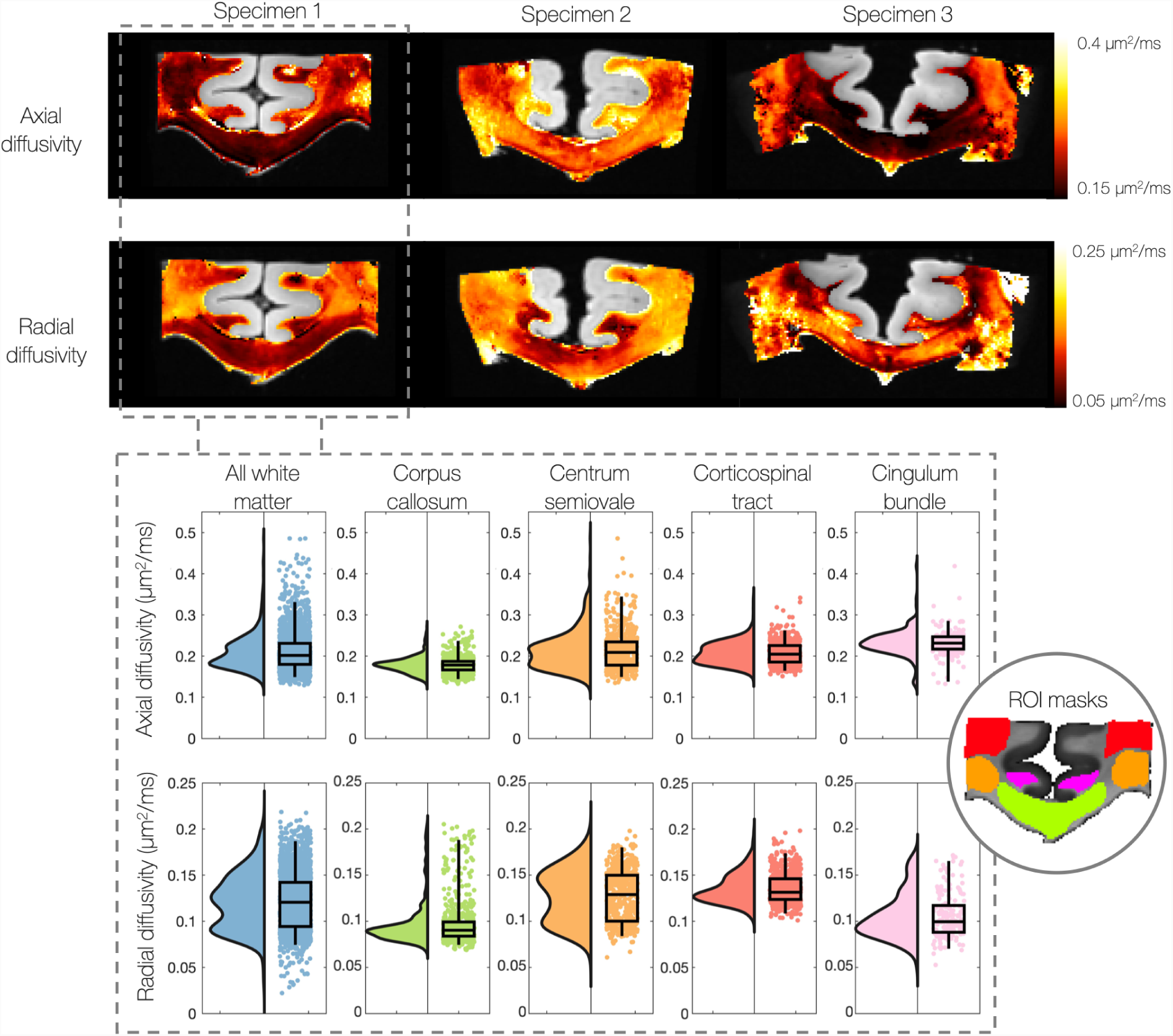
Heatmaps (top) and raincloud plots (bottom, specimen 1 only) show how the axial and radial diffusivities (and thus the FRF) vary on a voxel-wise basis across the white matter. We find notably lower radial diffusivities in the corpus callosum compared to other white matter tracts: the centrum semiovale, corticospinal tract and cingulum bundle. Additionally, we find two apparent distributions of radial diffusivities in the white matter. These results challenge standard CSD methods where a global FRF is assumed constant across the sample. Note the change in scale bar between the axial and radial diffusivities, with the radial diffusivity generally lower.

To assess how the diffusivities vary with anatomical regions of interest, Figure 7 (bottom) compares the distribution of axial and radial diffusion in all white matter voxels to those only in the corpus callosum, centrum semiovale, corticospinal tract and cingulum bundle. We see considerably lower diffusivities in the corpus callosum, as well as two distinct distributions of radial diffusivities across all white matter voxels.

### 3.4 Fibre response function

Figure 8 compares three ways of estimating the FRF from the dMRI data alone: the two conventional approaches estimate a global FRF, whilst the joint model estimates the FRF on a voxel-by-voxel basis. Using MRtrix3 [26], the FRF was first estimated from the 300 highest FA voxels and from the most-likely single-fibre voxels defined by the iterative Tournier algorithm [27]. This was compared to the joint model estimates of the FRF for voxels in the corpus callosum, centrum semiovale and corticospinal tract. To consider local changes in the FRF (rather than the voxel-wise variations of Figure 7), Figure 8 shows the mean FRF across a 2×2 neighbourhood of voxels in each anatomical region.

**Figure 8:**
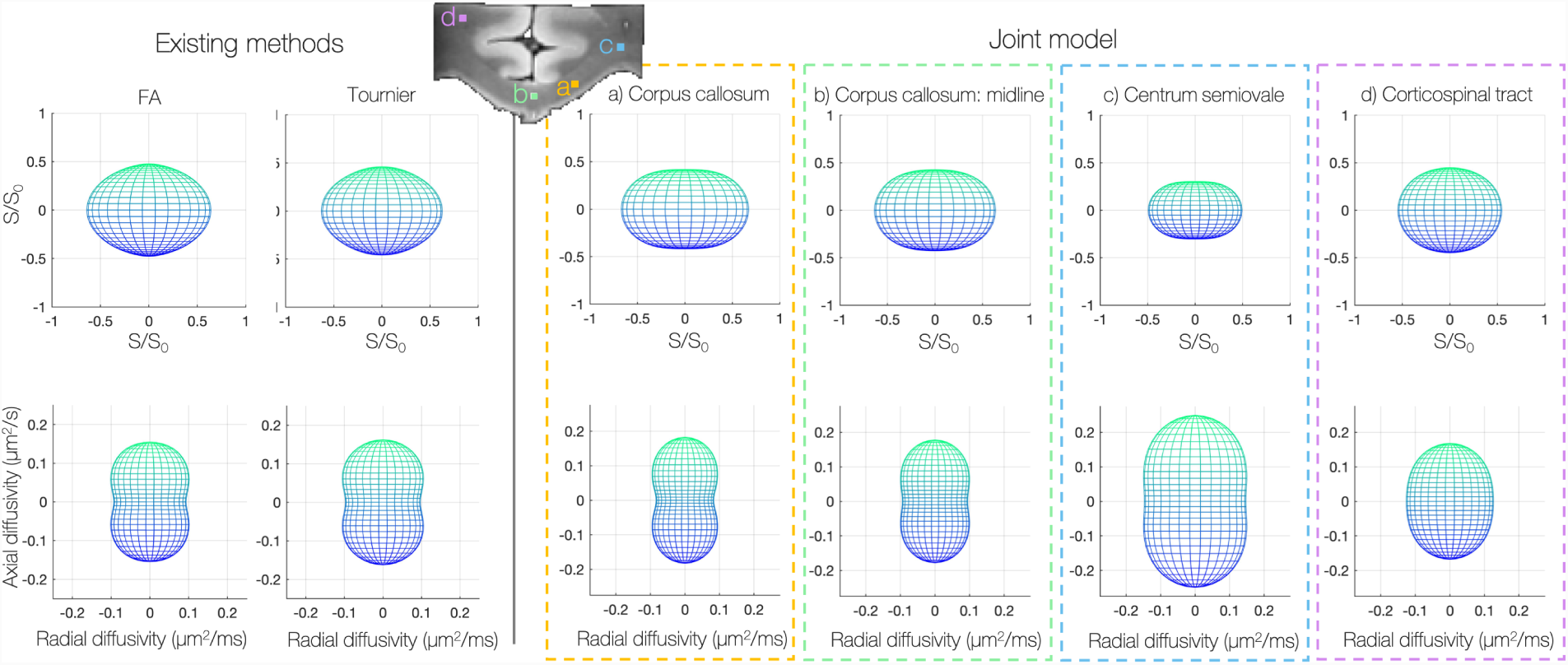
Variations in the observed fibre response function (FRF); here the FRF describes the diffusion profile of a single fibre bundle aligned vertically with respect to the page. The axial and radial diffusivities which describe the FRF (defined as an axially-symmetric diffusion tensor) are plotted below. A global FRF was first estimated from the 300 highest FA voxels and the Tournier algorithm [27] which selects the most likely ‘single-fibre’ voxels (left). This was compared to the FRF estimated by the joint model in the corpus callosum, centrum semiovale and corticospinal tract. The joint model estimates the FRF on a voxel-wise basis. However, here we consider local changes in the FRF; the FRFs shown were characterised by the mean axial and radial diffusivities over a 2×2 neighbourhood in each anatomical region. We see how the joint model predicts in general a more anisotropic (oblate) FRF in the corpus callosum and that the FRF appears to vary considerably with anatomy.

Both conventional methods typically estimate the FRF from voxels in the corpus callosum. Here the joint model estimated an FRF with lower radial diffusivity and higher diffusion anisotropy (*d*_*axial*_ - *d*_*radial*_) when compared to the conventional FRFs. This would be consistent with standard methods conflating fibre dispersion with diffusivity in the voxels used to generate the FRF.

In the joint model, the FRF appeared to vary considerably with neuroanatomy. Compared to the corpus callosum, Figure 8 depicts higher axial and radial diffusivities in the centrum semiovale and a highly spherical FRF in the corticospinal tract.

Notably, Figure 8 demonstrates FRF variability across the corpus callosum, which is typically considered a homogenous tract of coherently oriented fibres. Comparing the example FRF’s in Figure 8, the midline of the corpus callosum is characterised by lower axial diffusivity, higher radial diffusivity and a subsequent decrease in diffusion anisotropy.

### 3.5 FODs

Notable changes to the estimated FODs were also observed. Figure 9 (top) shows representative voxels from the corpus callosum of specimen 1 where, when compared to the CSD, the joint model estimated narrower FODs and a lower degree of dispersion in the underlying fibres. This finding was consistent across many (but not all) voxels in the corpus callosum of all three specimens.

**Figure 9:**
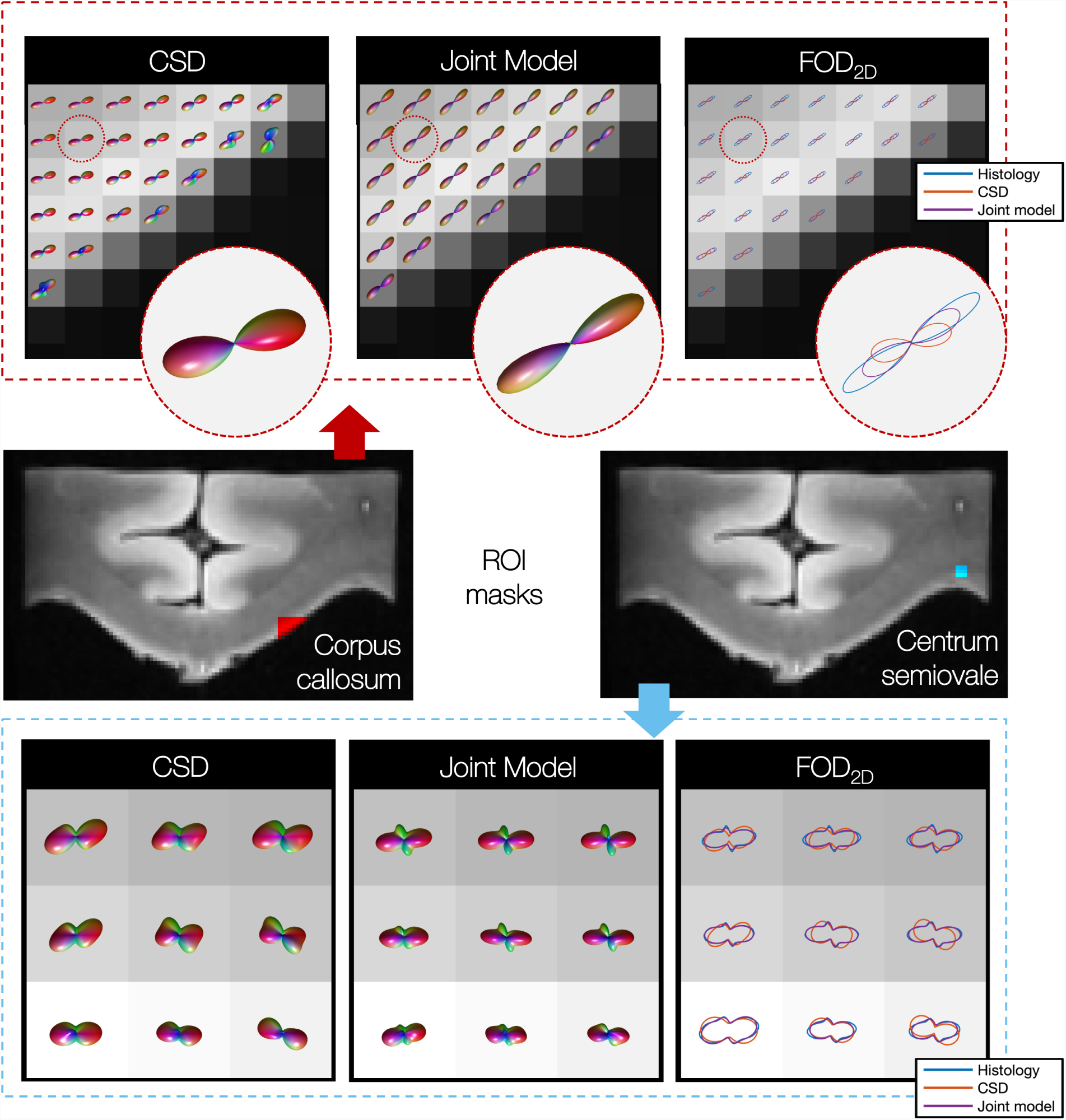
3D fibre orientation distributions (FODs) derived from CSD and the joint model in both the corpus callosum (top) and centrum semiovale (bottom). Top: The joint model estimates narrower FODs and lower dispersion in the underlying fibres. Bottom: In the centrum semiovale, additional distinct fibre populations are evident in the joint model FODs when compared to CSD. Notably, these peaks are recognisable in both the CSD FOD and the histology independently, and are not introduced by the histology only. In the right-hand pane the CSD and joint model FODs have been projected onto the plane of the histological data and are compared to the histology-derived FOD. Here we see the greatly improved fit of the joint model FOD to the histology.

Furthermore, in the joint model we sometimes observed the emergence of additional distinct fibre populations in the FOD. Figure 9 shows an example from the centrum semiovale; a known region of crossing fibres.

Figure 9 right (‘FOD_2*D*_’) shows the improved fit of the FOD when projected into the histological plane and compared to the histologically derived *FOD*_2*D,micro*_. This is quantified by the reduced KL divergence as shown in Figure 6. We see here how the histological data acts as a soft constraint on both the fibre orientation and amount of dispersion in the MRI voxel.

### 3.6 Limitations

Figure 10 demonstrates two cases where, due to limitations of the 2D histological data, the histology may bias the joint model prediction of the FOD. Firstly, the histology may underestimate, or totally omit, the volume fraction of a secondary fibre population (Figure 10 top). We see how the joint model thus attempts to minimise the presence of a secondary fibre population, the resultant FOD being perhaps biased or biologically improbable.

**Figure 10:**
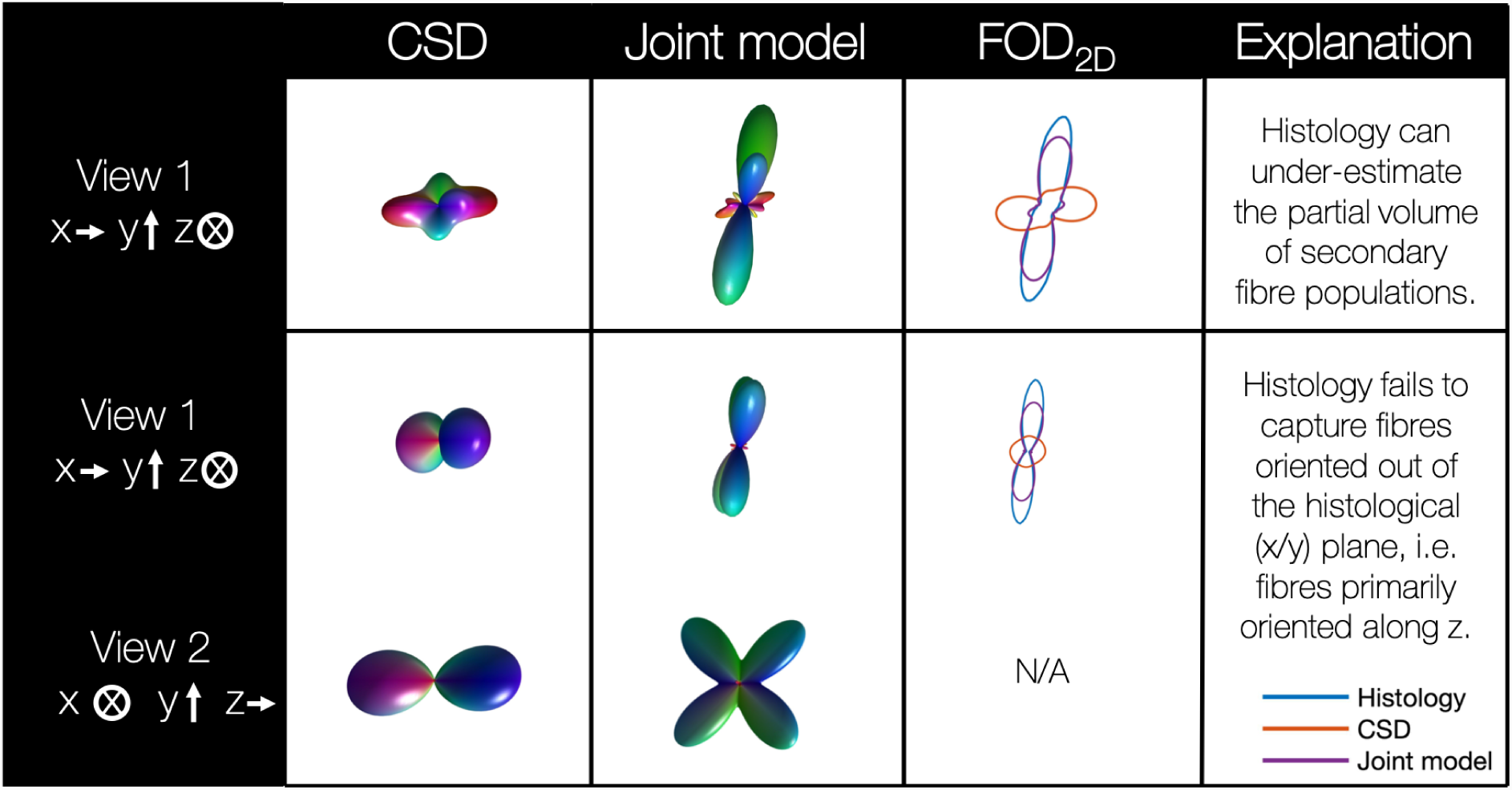
Two cases where the histological data might bias the 3D FOD predicted by the joint model. Histology may under-estimate the volume fraction of in-plane secondary fibre populations (top), or omit out-of-plane fibres from the histology-derived 2D FOD (bottom). Here the 3D FOD is seen from two orthogonal views: looking onto (view 1) and out of (view 2) the histological plane (denoted x/y).

Secondly, the histological data is most sensitive to fibre bundles in, or close to, the histological plane. Figure 10 (bottom) shows a single voxel from two orthogonal views: looking down on the histological plane (view 1), and perpendicular to the plane (view 2). In CSD, we see a single fibre population, primarily oriented out of the histological plane; the CSD-derived 2D FOD is consequently highly spherical. The histology-derived FOD however looks substantially different. This may be due to the histology-derived FOD over-emphasising either in-plane components of mostly through-plane fibres or minor fibre populations which lie in the histological plane. Alternatively, there could be mis-registration of the dMRI and histology data such that the two datasets are incompatible. Consequently, as the joint model tries to fit the histological data, it appears to split the single-fibre FOD in two. Both examples in Figure 10 were taken from the centrum semiovale on the left hand side of specimen one: a known region of complex fibre configurations and, in this specimen, a region of high dMRI-associated error in the joint model.

### 3.7 Orientation of the FOD peak

In the joint model, the histological data constrains both the fibre orientation and amount of dispersion in each dMRI voxel. This is likely to affect the orientation of the estimated FOD peak in two respects. Here we define the peak orientation by the azimuthal angle in, and inclination out of, the histological plane. Firstly, any mis-registration of the dMRI and histology data may cause a rotation in the azimuthal angle. Secondly, as the histological data is most informative of in-plane fibres, the joint model may favour FODs of low inclination. Figure 11 investigates these effects by comparing the orientation of the primary peaks of FODs estimated by both CSD and the joint model. Figure 11a shows the angular deviation of the two peaks where the three samples have a median difference of 12.4°, 11.2° and 10.7°. Figures 11b and c compare the azimuth and inclination of the primary peaks respectively. We see how all three specimen show very close correlation with the robust regression line close to the line of unity and r-values between *r* = 0.97 - 0.99. Note that here we use robust linear regression to limit our sensitivity to outliers and that the r-values of robust regression (here calculated using MATLABs *fitlm* [30]) are typically inflated when compared to standard linear regression.

**Figure 11:**
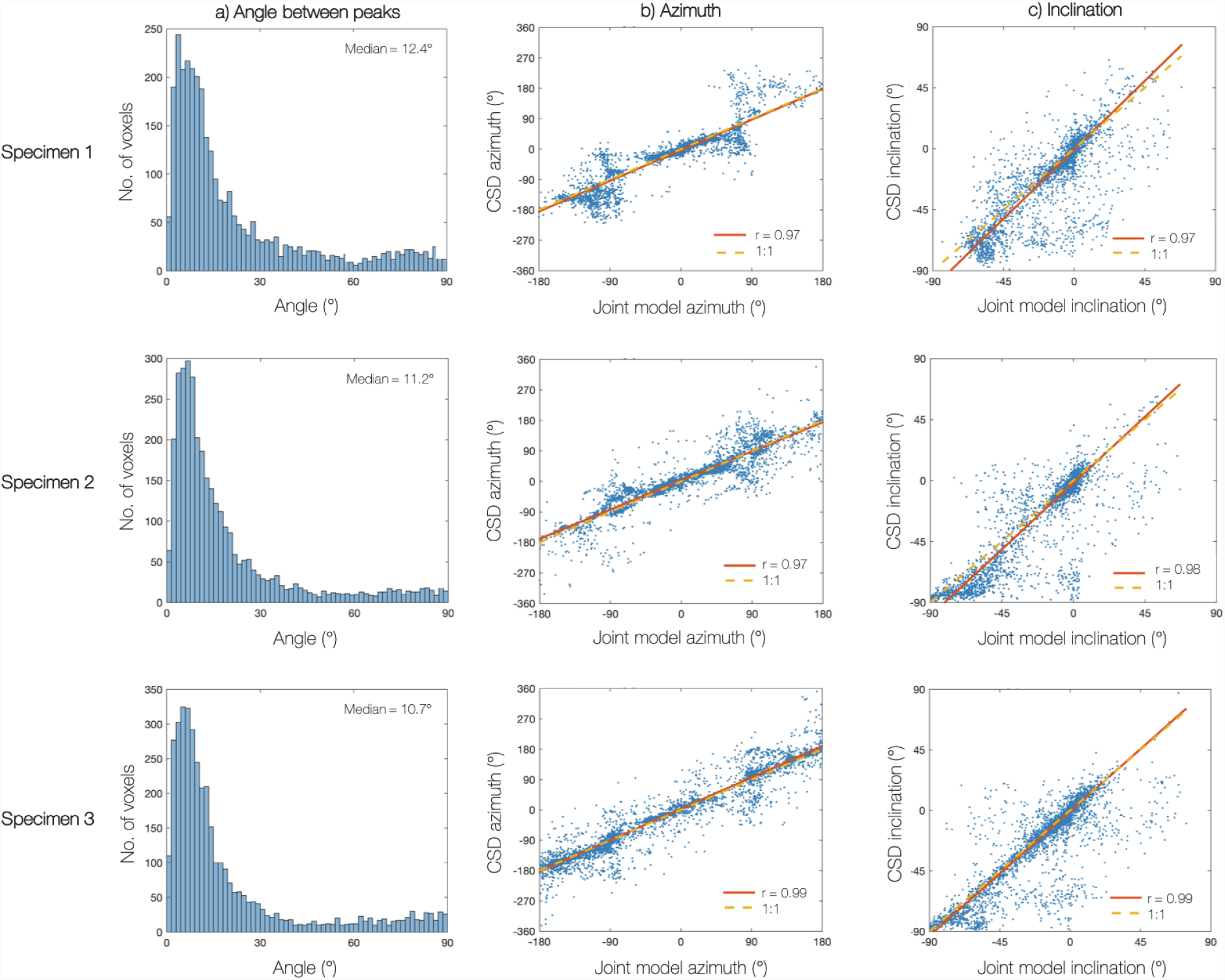
Comparing the orientation of the primary FOD peak derived from both CSD and the joint model. The peak orientation is described by an azimuth and an inclination angle which is defined with respect to the histological plane (inclination = 0°). In the joint model, any mis-registration of the dMRI and histology data may result in a rotation of the peak azimuth. In addition, as histology is most informative of fibres lying in or close to the histological plane, the joint model might favour FODs of low inclination angles. In all 3 specimen there is a low angular deviation (a) between the CSD and joint model FOD peaks, with good correlation between the azimuthal (b) and inclination (c) angles: the robust regression lines are close to unity with r-values between 0.97-0.99.

In 11b specimen 1 we see a prevalence of joint model estimated FOD peaks at azimuthal angles of *θ* ± 90°, i.e along the inferior-posterior axis. This is due to an almost equal prevalence of histology-defined FODs with peaks at these angles and is probably caused by an artefact in the histology data. This however does not appear to greatly impact either the inclination (Figure 11c) or total angular difference (Figure 11a) between the CSD and joint model derived FODs of specimen 1. Overall, Figure 11 indicates a markedly stable relationship between the ODF peaks derived from CSD (dMRI analysis only) and the joint model (combined dMRI and histology analysis).

## 4. Discussion

In the joint model we utilised a data-fusion framework to combine co-registered dMRI and histology data from the same tissue samples, acquired from three postmortem human brains. This allowed us to test the validity of a brain-wide fibre response function as used in CSD [16]. With the inclusion of histological data, we were able to constrain both the fibre orientation and amount of dispersion within each MRI voxel, albeit in 2D. Consequently, the joint model could overcome the degeneracy between fibre dispersion and radial diffusion to estimate the FRF and FOD simultaneously on a voxel-by-voxel basis.

Using simulated data, we evaluated the model specificity and accuracy. For simulated data of single-fibre populations, the joint model provided particularly robust estimates of both orientation dispersion and radial diffusion for fibre populations in or close to the histology plane (up to inclinations of 60°), where the histological data was most informative. However, in both simulation and post-mortem data the joint model was less effective at reconstructing out-of-plane fibre populations. This was due to an inherent limitation of the microscopy technique, where fibres oriented out of the plane aren’t readily detected in the 2D histological data. The bias of the model could be reduced if the microscopy was able to reconstruct the FOD in 3D; 3D polarised light imaging developed by Axer *et al.* [37], or 3D confocal microscopy from Schilling *et al.* [38, 23] being two examples of such techniques. In addition, future work could take a more heuristic approach where the regularisation factor *λ*_*micro*_ could depend on fibre inclination, as a proxy of how informative the microscopy data is of the 3D FOD.

In this joint model, the FRF was characterised by axial and radial diffusivities, which appear to vary considerably across the human brain. Generally, both diffusivities seem to follow a similar pattern of variation, with particularly low values of radial diffusivity found in the corpus callosum. Contrary to the assumption of a brain-wide FRF in CSD, our results demonstrate that the FRF is dependent on the local anatomy. This supports recent findings by Schilling et al. who used 3D confocal microscopy to investigate FRF variation across the brain of a squirrel monkey [23]. FRF variation is expected as tissue characteristics such as axonal density, packing and myelination are unlikely to hold across the entire brain [20]. For example, areas of the corpus callosum have been reported to have only 30% myelination [39] which would significantly impact the FRF; decreased myelination has been shown to be associated with high radial diffusivity [40, 41, 42]. Indeed, characteristically different fibres should each have their own unique FRF, although in practice this would be difficult to model.

Not only does FRF variation challenge the assumption of the brain-wide FRF used in CSD, but also implies that current analyses may be biasing the shape of their reconstructed FODs throughout the brain. Indeed, in known regions of crossing fibres such as the centrum semiovale, the joint model estimated distinct secondary fibre populations which were often recognisable but not well defined in the CSD-derived FODs. Furthermore, the joint model often estimated FODs of lower dispersion in single-fibre voxels, particularly across the corpus callosum. Current algorithms which estimate the FRF directly from the data [17, 27, 27] typically estimate the FRF from voxels in the corpus callosum which exhibit high fractional anisotropy. However, even these voxels are unlikely to contain a perfectly coherent fibre bundle. Indeed, recent studies of both histology [43], polarised light imaging and dMRI data [10] have found significant levels of dispersion in the corpus callosum, particularly at the midline [10]. To obtain accurate estimates of the FOD, both this study and that by Schilling *et al.* [23] support the development of analyses which estimate a local rather than global FRF. Both approaches are however limited as they require co-registered dMRI and microscopy data in postmortem tissue samples. Building on the work by Lasič *et al.* [13] and Szczepankiewicz et al. [14], Cottaar *et al.* combine linear and spherical tensor diffusion encoding to overcome the degeneracy between fibre dispersion and diffusion anisotropy to estimate both on a voxel-wise basis, with highly consistent results. Similar techniques could provide exciting new avenues for local FRF estimation in *in vivo* data and more faithful FOD reconstruction in spherical-deconvolution based analyses.

Multimodal datasets also present a number of challenges: one being that the microscopy data is only representative of a subset of the tissue microstructure. Here the tissue sections were stained to visualise the myelin content of the tissue, yet only pixels above a certain staining intensity were recognised as myelinated fibres. Fibres that failed to meet the staining threshold (perhaps those with with less myelin and typically lower diameter), and other cell types (e.g. glia) may have therefore been underrepresented or omitted from the microscopy-derived FOD. Secondly, as the histological stain was only sensitive to myelin, other cell types (e.g. glia) were excluded from the microscopy FOD. If the unmyelinated axons or glia both contributed to the diffusion signal and acted to increase the dispersion of the true FOD, the FOD of the joint model would underestimate the amount of dispersion compared to that of the true contributing microstructure. Nonetheless, as seen in Figure 9, the joint model did not act to fit to the histological data perfectly. Indeed the joint model often estimated a slightly higher degree of dispersion than prescribed by the histology and on occasion assumed a substantially different FOD, if this was both driven and supported by the dMRI data. This is a result of the histology acting as a soft constraint, rather than fixing the fibre orientation and dispersion.

The postmortem dMRI data used here presents a second challenge. Although the data provides high spatial resolution (0.4 mm isotropic), it has relatively low diffusion contrast. This is likely due to the postmortem interval (time between death and fixation) and immersion fixation of the human brains from which our samples were taken. It is most likely that the meninges, specifically the pia mater and the components of the dura, were attached during immersion fixation of the postmortem brains in this study. As the samples originated from the centre of each brain, the uptake of fixative to this area will have been particularly slow as fixative will have had to diffuse from the cortical surface to reach the callosum, increasing the apparent postmortem interval. Prior to fixation, post-mortem tissue may begin to decompose, wherein both the postmortem interval and fixation method have been shown to change the diffusion properties of the tissue [44, 45]. This likely explains both the inter-specimen variation in estimated diffusivities and why the diffusivities reported here are low when compared to *in vivo* data where, for comparison, the NODDI model assumes *d*_*axial*_ = 1.7 *µm*^2^*/ms* [46, 32]. It is important to note that the joint model generalises to any dataset which includes dMRI alongside microscopy data from which we can estimate fibre orientations. This encompasses existing dMRI and histology datasets [38, 47, 10], as well as data from alternative microscopy techniques such as the mesoscopic fibre orientations obtained from polarised light imaging [37, 8, 10]. As such, future work aims to apply the CSD-based joint model to other datasets, of both human and monkey (perfusion-fixed at death) brain tissue.

Finally, this report looks through the lens of a convolution-based joint model to detail both the advantages and challenges of a dMRI and microscopy data-fusion framework. As such, we recognise this model to be only one example of a joint modelling approach. The microstructure model at the centre of the data-fusion framework can itself take a range of forms, or be extended to increase both its complexity and specificity. For example, future work will consider a multi-compartment model of the fibre response function, which distinguishes intra-from extra-axonal space, each characterised by a unique diffusion profile. In this manner, we hope to disentangle which tissue compartment is driving the FRF variation across the brain.

## 5. Conclusion

The joint model takes a data-fusion approach, combining dMRI and histology data from the same tissue sample to investigate the diffusion properties of white matter under conditions where multiple modalities are informative of the ‘true’ tissue microstructure. In the model, histology acted as a soft constraint on both the fibre orientation and amount of dispersion in each dMRI voxel. This allowed us to overcome the degeneracy between fibre orientation dispersion and radial diffusion, to estimate the FOD and FRF on a voxel-by-voxel basis. Our results demonstrate how the diffusion characteristics of a single fibre (here characterised by axial and radial diffusivities) vary considerably across the brain. Notably, the diffusivities were found to vary both between and within white-matter tracts; even in the corpus callosum where the microstructure is typically considered fairly homogenous. These findings contradict the assumption of a brain-wide FRF, currently used in many deconvolution-based analyses. Finally, our results suggest that current diffusion models may be overestimating dispersion in single-fibre voxels and underestimating the number of distinct fibre populations in known regions of crossing fibres; both of which have important implications for tractography [48].

## 6. Acknowledgements

This work was supported by the Wellcome Trust (grant WT202788/Z/16/A), EPSRC and MRC (grants EP/L016052/1 and MR/L009013/1). The Wellcome Centre for Integrative Neuroimaging is supported by core funding from the Wellcome Trust (203139/Z/16/Z). We acknowledge the Oxford Brain Bank, supported by the Medical Research Council (MRC), the NIHR Oxford Biomedical Research Centre and the Brains for Dementia Research programme, jointly funded by Alzheimers Research UK and Alzheimers Society for supplying the postmortem brains. We also thank the donors and their families.

